# The *gac* system integrates physical and chemical cues to promote plant root attachment

**DOI:** 10.64898/2026.04.10.717875

**Authors:** Guy Sobol, David M. Hershey

## Abstract

Plants host complex communities of microbes that are attached to root surfaces. While many studies have sampled mutant populations after prolonged incubation on roots to identify bacterial genes that enable long-term colonization, the molecular mechanisms governing the early stages root attachment remain less understood. Here, we developed an *in vitro* root culture system that enables controlled and scalable investigation of bacterial attachment to root tissue. We used this platform to perform a genome-wide screen for root attachment determinants in the plant-associated bacterium *Pseudomonas protegens* Pf-5. Our results reveal that the *gacSA* two-component system functions as a sensory integration hub for coordinating early root attachment. Mutations that disrupt *gacS* or *gacA* cause severe root attachment defects despite having no effect on abiotic surface attachment in standard biofilm assays. Mutation of flagellar assembly genes enhances root attachment by mimicking surface contact and activating the *gac* system. In parallel, chemical cues released by roots stimulate surface attachment in a *gac* dependent manner. By integrating these signals, the *gac* system activates cyclic di-GMP-mediated attachment programs that drive the transition from planktonic to sessile behavior required for root association. We build on this model to show that manipulating flagellar surface sensing enhances the competitive fitness of Pf-5 in the presence of a synthetic bacterial community, suggesting a strategy to improve the competitive fitness of beneficial microbes on crops. These findings establish a mechanistic framework linking surface sensing, global regulation, and root attachment in a beneficial rhizobacterium.

**Importance:** Plant roots are covered with diverse microbes that strongly influence plant health. Growing in association with roots has many benefits, but how bacteria attach to root tissue remains poorly understood. We developed a system to study how a bacterium that improves plant growth called *Pseudomonas protegens* Pf-5 attaches to root tissue. We found that physical contact with the root surface and chemical cues released by roots both enhance attachment to root tissue. A sensory system called *gacSA* is responsible for integrating physical and chemical cues to activate a root attachment program. Variant bacteria that prematurely activate the *gac* system compete more effectively with other bacteria on roots, suggesting that the root attachment pathway we characterized could serve as a strategy to use beneficial bacteria in agriculture.

## Introduction

Surface-associated communities called biofilms represent the predominant lifestyle for most bacteria (1, 2). Cells in biofilms are encased in a self-produced extracellular matrix that provides resistance to stresses such as shear flow, unfavorable temperatures, antimicrobial substances, and nutrient limitation (3). While biofilms are found in nearly all environments on earth, their association with living tissues has particularly significant effects on humans. Biofilms within human tissues are thought be the source of chronic infections, and tolerance to antimicrobial treatments makes them difficult to treat in clinical settings (4). In agricultural systems, biofilms that from on plant tissues are crucial determinates of crop productivity (5). Despite the profound importance of host-associated biofilms, mechanistic analyses of biofilm formation have historically been dominated by studies on abiotic surfaces such as plastic, glass, and agar. Knowledge of host-associated biofilms is often inferred from these inanimate models (6). Increasing efforts to study bacterial colonization of biotic substrates have begun to reveal substantial differences in host-associated biofilm formation, but continued work is needed to accelerate the development of strategies for managing biofilms in host-associated settings (7).

Plant health is strongly influenced by complex biofilms in the rhizosphere, a narrow zone of soil surrounding root tissues (8–10). Plants interact with microbes in the rhizosphere by secreting sugars, amino acids, organic acids, and secondary metabolites that are collectively referred to as exudates (11, 12). Exudates create chemical gradients that attract bacteria toward the root surface (Fig. 1). They also promote bacterial aggregation and provide nutrients that make the root surface a favorable niche (13). Plants, in turn, rely on microbes in the rhizosphere for nutrient acquisition, suppression of pathogenic organisms, and tolerance to stressors such as drought. This mutualistic relationship requires microbes to physically attach to the root surface.

**Figure 1.**
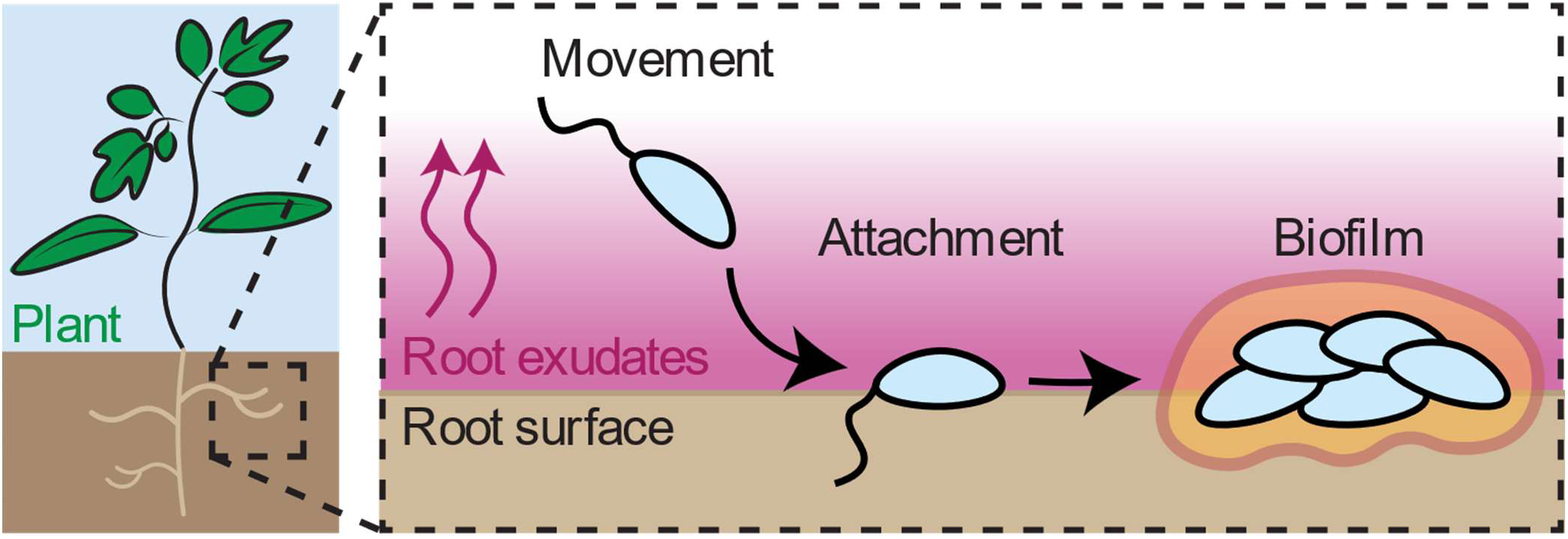
Bacterial attachment to root surfaces Roots release a mixture of exudates into the surrounding soil, generating a concentration gradient that is highest near the root surface (illustrated as a pink gradient). Rhizosphere bacteria actively move toward roots based on the exudate gradient, attach to the root surface via secreted adhesins, and subsequently form a biofilm.

**Figure 2.**
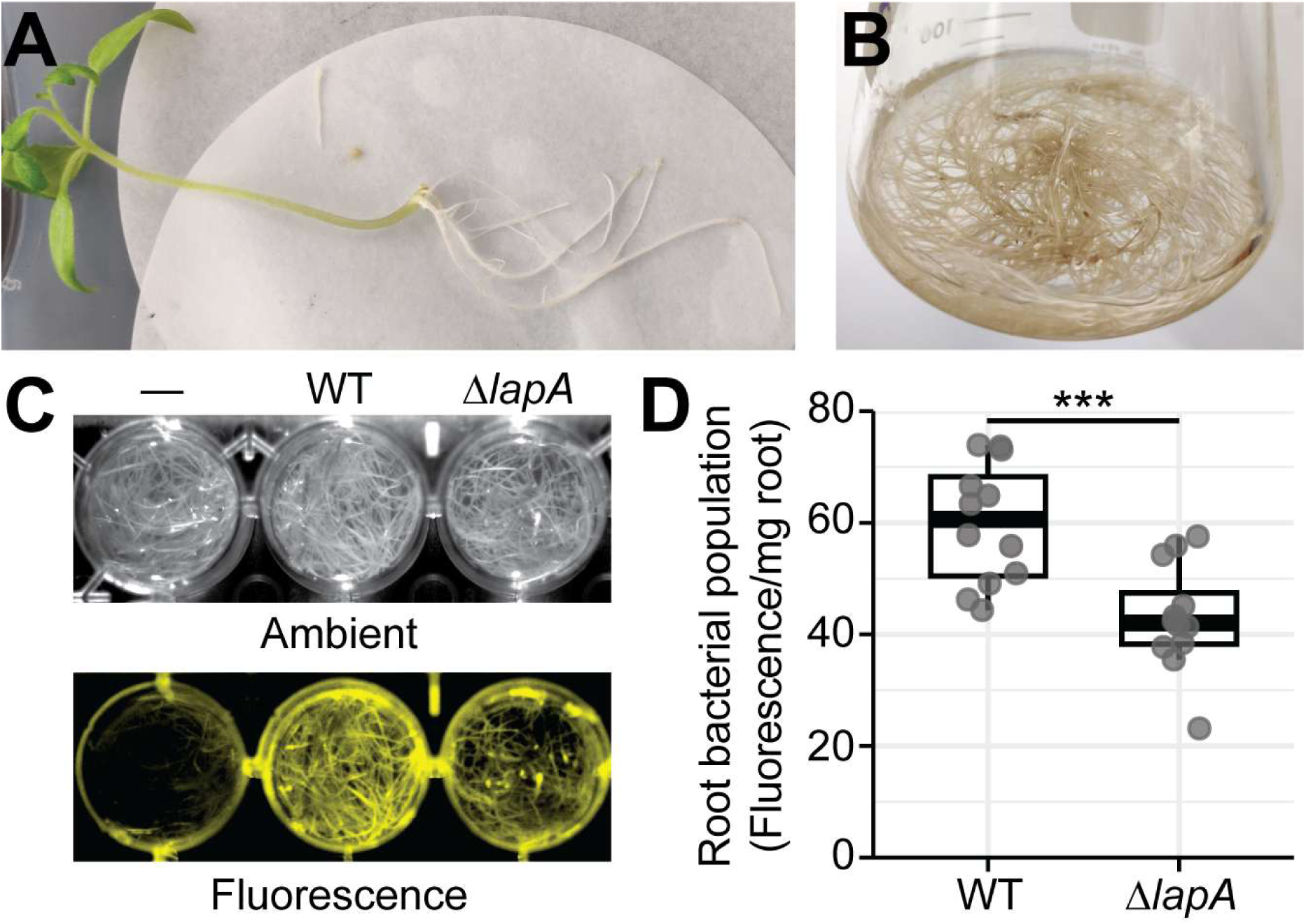
Establishing an *in vitro* root tissue culture system to study root colonization (A) Seedlings grown under sterile tissue culture conditions were inoculated with *Agrobacterium* at an incision at the base of the hypocotyl to generate hairy roots. (B) Excised root tissue transferred to liquid medium and propagated in shaking culture, resulting in root biomass suitable for downstream colonization assays. C and D, Root tissue placed in plate wells was inoculated with a bacterial suspension of mVenus-tagged Pf-5 wild-type or Δ*lapA*. Root colonization was evaluated by measuring fluorescence from roots (C) or quantified by recovering the root-attached bacterial fraction and measuring its emitted fluorescence (D). Bold lines within each box represent median values. Top and bottom sides of the boxes represent the third and first quartiles of the value distribution, respectively. Lines extending from the boxes denote the extreme values within 1.5 times the interquartile range. Statistical significance was determined by a pairwise *t*-test. Asterisks indicate *P* < 0.001. Data are pooled from three independent experiments.

Efforts to identify the genetic determinants of root attachment have largely relied on inoculating mutant libraries onto plants growing in substrates such as soil, vermiculite, sand, or synthetic plant media. Sampling the root-associated population using transposon sequencing (Tn-Seq) has consistently identified genes involved in amino acid biosynthesis, carbon utilization, nitrogen assimilation, iron acquisition, suppression of plant defenses, and competition with other microbes (14–23). Such studies have been instrumental in defining metabolic pathways required for persistence in the rhizosphere. However, the prolonged incubations emphasize sustained colonization rather than the earliest stage of root attachment. A more thorough understanding of how bacteria attach to root tissue is needed to understand biofilm formation in the rhizosphere.

Members of the *Pseudomonas fluorescens* species complex are frequent residents of the rhizosphere (24, 25). These bacteria often promote plant health by enhancing nutrient uptake, producing growth-promoting compounds, and antagonizing pathogens. There is extensive knowledge of the mechanisms that govern biofilm formation within the *Pseudomonas* genus, particularly in the opportunistic human pathogen *Pseudomonas aeruginosa* (26). *Pseudomonads* coordinate biofilm formation through a regulatory paradigm centered on the second messenger cyclic-di-GMP (c-di-GMP). While the details of each signaling cascade vary among individual strains, environmental cues trigger elevated c-di-GMP levels which in turn bind effector proteins that promote biofilm-associated behaviors such as reduced motility, biofilm matrix production, and expression of adhesins. A well-characterized output of c-di-GMP signaling in *P*. *fluorescens* is the Lap system, which controls a large cell-surface adhesin LapA. Elevated intracellular c-di-GMP concentrations promote the accumulation of LapA at the cell surface where it mediates attachment and supports biofilm formation. Reduced c-di-GMP levels result in LapA cleavage and release, leading to detachment and dispersal (27).

In this study, we develop an *in vitro* root tissue culture system to investigate bacterial root attachment. This approach overcomes practical limitations of studying root attachment in whole plants and avoids abiotic adhesion assays that do not capture interactions with living host tissue. We use *Pseudomonas protegens* Pf-5, a soil bacterium that associates with the roots of various crops (28–32), to perform a genome-wide screen for root attachment determinants. Our results demonstrate that the *gacSA* two-component system acts as a key integrator of signals to coordinate root attachment. Both flagellum-mediated surface sensing and chemical cues released by roots activate *gac* signaling to promote c-di-GMP-dependent attachment. Furthermore, we show that manipulating flagellar surface sensing enhances the competitive fitness of Pf-5 in the presence of a synthetic bacterial community, suggesting a potential strategy for improving the performance of beneficial bacteria in agricultural applications. Our results shed new light on how host tissues influence the earliest stages of biofilm formation.

## Results

### Genome-wide screen identifies *Pseudomonas protegens* Pf-5 genetic determinants for early root attachment

We developed an *in vitro* tissue culture system for propagating tomato roots in liquid medium (Fig. 1A, B). This platform allows large volumes of root tissue with an extensive surface for attachment to be cultivated independently of whole plants under highly controlled conditions. We then engineered a Pf-5 strain expressing mVenus and used it to carefully optimize conditions for robust colonization of root tissue. Pf-5 was inviable in standard root culture medium, but adding carbon sources commonly detected in root exudates allowed Pf-5 to grow (Fig. S1A-C) (33). We validated our system by comparing root association of fluorescently-tagged wild-type Pf-5 to a mutant lacking the large surface adhesin LapA (27). Bacterial abundance on roots was measured by imaging total emitted fluorescence from roots in plate wells (Fig. 1C), measuring the fluorescence emitted by cells recovered from roots or directly quantifying viable cell counts (Fig. 1D; Fig. S1D). Both strains attached to root tissue, but roots inoculated with the Δ*lapA* mutant emitted reduced levels of fluorescence and carried less CFUs compared with wild-type Pf-5, consistent with a deficiency in root attachment.

We used the *in vitro* colonization system to perform a genome-wide screen for Pf-5 genes involved in attachment to roots. A barcoded transposon mutant library was inoculated into flasks containing root tissue, and the mutant pool was allowed to attach to roots for 24 hours (34). We predicted that mutants that retained the ability to associate with root tissue would partition out of the medium as they attached to roots, enriching for attachment deficient mutants in the liquid medium. We amplified these selections by inoculating the unattached population into fresh root tissue for four additional passages (Fig. 3A). Barcode abundances from the initial mutant population and from the unattached fraction after each root passage were used to calculate fitness scores for each gene across the passages (Fig. 3B; Table S1).

**Figure 3.**
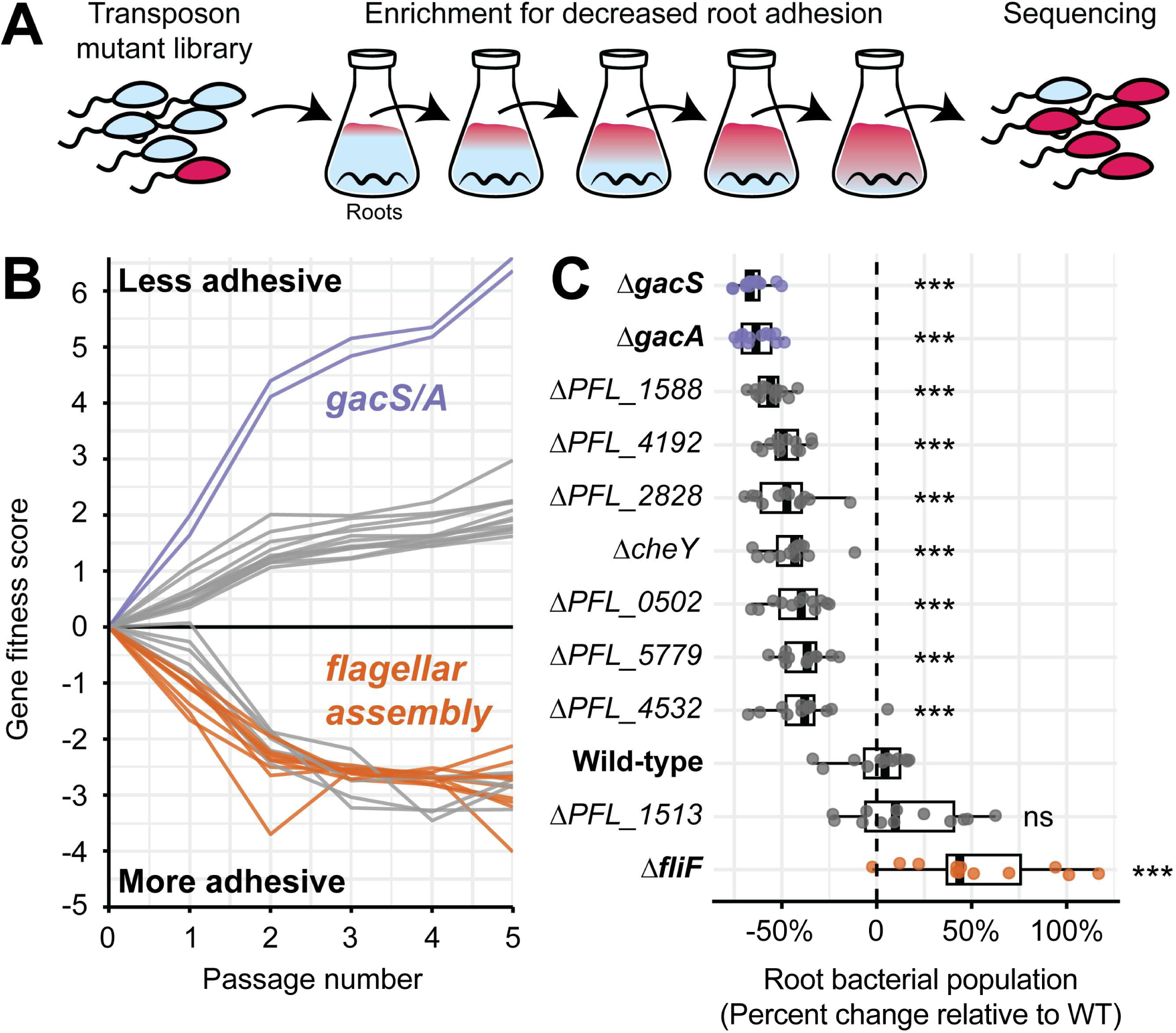
A genome-wide screen identifies genetic determinants for root attachment (A) A pooled transposon mutant library carrying mutants impaired in root association (red) was inoculated into flasks containing hairy roots. After incubation, the unattached (planktonic) fraction was collected and serially passaged into fresh root-containing flasks for five rounds. The resulting population is enriched for adhesion-defective mutants and sequenced. (B) Fitness trajectories for the 15 most hypo-and 15 most hyper-adhesive mutants identified in the root passaging Tn-Seq screen. Lines connect fitness scores for each gene across passages 0-5. Positive fitness scores indicate mutants that increased in abundance in the unattached (media) fraction, consistent with reduced root adhesion, whereas negative scores reflect depletion from the media and therefore enhanced adhesion to roots. Genes highlighted in purple represent the *gacS* and *gacA* genes, orange denotes flagellar assembly-associated genes, and gray indicates genes outside these categories. (C) Root colonization phenotypes for in-frame deletion mutants of genes identified in the screen. Root colonization was quantified by recovering the root-attached fraction of cells and measuring its emitted fluorescence. Bold lines within each box represent median values. Top and bottom sides of the boxes represent the third and first quartiles of the value distribution, respectively. Lines extending from the boxes denote the extreme values within 1.5 times the interquartile range. Statistical significance was determined by a pairwise *t*-test between wild-type and each strain. Asterisks indicate *P* < 0.001, “ns” no significant difference. Data are pooled from three independent experiments.

Genes with the strongest effects on root attachment are shown in Tables 1 and 2. Mutations causing the strongest defects in root attachment mapped to *gacS* and *gacA* which encode a histidine kinase-response regulator pair that sits atop a large regulatory hierarchy. Mutations affecting transcription factors (*PFL_2828*, *PFL_0502*, *PFL_1588*), uncharacterized transporters (*PFL_4192*, *PFL_1740*), the *pxpABC* operon encoding the allophanate hydrolase enzyme (*PFL_1513*-*1515*) (35), as well as genes involved in c-di-GMP (*PFL_5779*, *PFL_4532*), lipid (*PFL_5687*), and carbon (*PFL_6157*) metabolism also caused root attachment deficiencies.

**Table 1:**
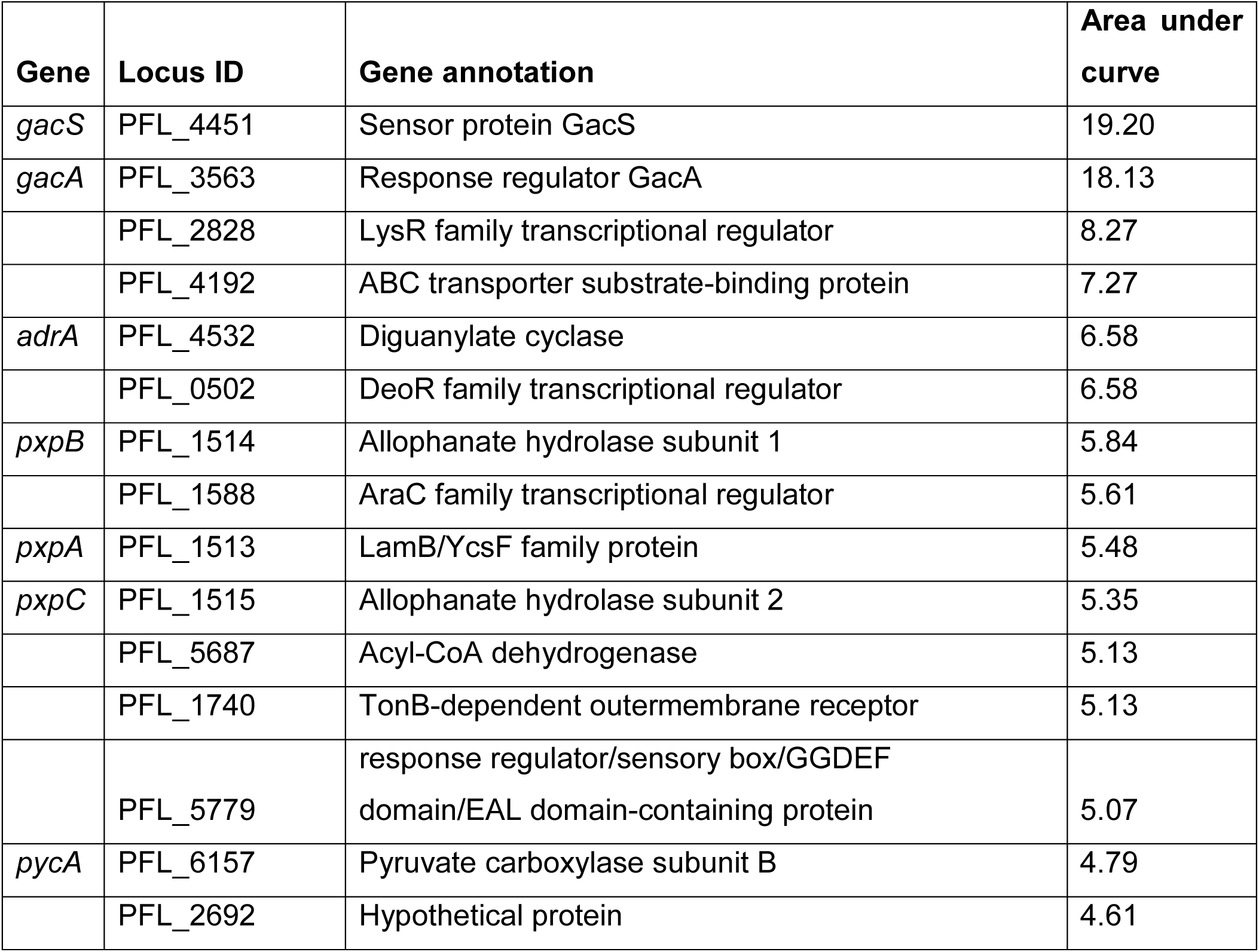
Top 15 protein-encoding hits enriched in the non-root-associated fraction (reduced attachment).

| Gene | Locus ID | Gene annotation | Area under curve |
| --- | --- | --- | --- |
| *gacS* | PFL_4451 | Sensor protein GacS | 19.20 |
| *gacA* | PFL_3563 | Response regulator GacA | 18.13 |
|  | PFL_2828 | LysR family transcriptional regulator | 8.27 |
|  | PFL_4192 | ABC transporter substrate-binding protein | 7.27 |
| *adrA* | PFL_4532 | Diguanylate cyclase | 6.58 |
|  | PFL_0502 | DeoR family transcriptional regulator | 6.58 |
| *pxpB* | PFL_1514 | Allophanate hydrolase subunit 1 | 5.84 |
|  | PFL_1588 | AraC family transcriptional regulator | 5.61 |
| *pxpA* | PFL_1513 | LamB/YcsF family protein | 5.48 |
| *pxpC* | PFL_1515 | Allophanate hydrolase subunit 2 | 5.35 |
|  | PFL_5687 | Acyl-CoA dehydrogenase | 5.13 |
|  | PFL_1740 | TonB-dependent outermembrane receptor | 5.13 |
|  | PFL_5779 | response regulator/sensory box/GGDEF domain/EAL domain-containing protein | 5.07 |
| *pycA* | PFL_6157 | Pyruvate carboxylase subunit B | 4.79 |
|  | PFL_2692 | Hypothetical protein | 4.61 |

**Table 2:** Top 15 protein encoding hits depleted from the non-root-associated fraction (enhanced attachment).

| **Gene** | **Locus ID** | **Gene annotation** | **Area under curve** |
| --- | --- | --- | --- |
| *fleE* | PFL_1637 | Flagellar hook-basal body complex protein FliE | 11.63 |
| *cheA* | PFL_1670 | Chemotaxis protein CheA | 10.58 |
|  | PFL_2045 | Amino acid aminotransferase | 9.93 |
| *fliF* | PFL_1638 | Flagellar basal-body MS-ring/collar protein FliF | 9.81 |
| *flgF* | PFL_1613 | Flagellar basal-body rod protein FlgF | 9.74 |
| *fliI* | PFL_1641 | Flagellar protein export ATPase FliI | 9.64 |
| *fliG* | PFL_1639 | Flagellar motor switch protein FliG | 9.53 |
| *fliA* | PFL_1667 | RNA polymerase sigma factor FliA | 9.43 |
| *flgE* | PFL_4477 | Flagellar hook protein FlgE | 9.43 |
| *fliK* | PFL_1646 | Flagellar hook-length control protein FliK | 9.42 |
| *flgG* | PFL_1614 | Flagellar basal-body rod protein FlgG | 9.39 |
| *flgC* | PFL_4479 | Flagellar basal body rod protein FlgC | 9.16 |
| *cheY* | PFL_1668 | Chemotaxis response regulator CheY | 9.13 |
| *fleR* | PFL_1636 | Sigma-54 dependent transcriptional regulator | 9.11 |
|  | PFL_1752 | HAMP domain-containing sensor histidine kinase | 8.85 |

Mutations that enhanced root attachment were concentrated in genes required for the function of the flagellum, including assembly regulators, structural components and chemotaxis systems (36). In-frame deletion mutants Δ*gacS* and Δ*gacA* recapitulated the strong attachment defect, with root attachment reduced by ∼60% relative to wild-type (Fig. 3C). The Δ*fliF* mutant, representing flagellar structural mutants and lacking a flagellum, attached ∼30% more effectively than wild type. Deletion mutants of *PFL_1588*, *PFL_4192*, *PFL_2828*, *PFL_0502*, *PFL_5779*, and *PFL_4532* were hypoadhesive, consistent with the Tn-Seq results. The Δ*cheY* mutant was hypoadhesive despite appearing hyperadhesive in the Tn-Seq results. Δ*PFL_1513* did not differ significantly from wild type, although it was hypoadhesive in the Tn-Seq results. All the deletion mutants displayed comparable growth kinetics to wild type (Fig. S2A).

### Flagellar-dependent surface sensing activates the *gac* system

We sought to understand two groups of genes with opposing effects on root colonization. *gacS* and *gacA* mutants have severe root attachment defects (Fig. 3). Activation of the *gacSA* two-component system induces expression of small RNAs called *rsm,* which go on to activate genes involved in biofilm formation, secondary metabolite production, and exopolysaccharide synthesis (37). Mutating flagellar genes in Pf-5 strongly enhances root attachment (Fig. 3). The flagellum plays numerous roles throughout the process of biofilm formation including facilitating productive contact with surfaces and recognizing physical cues for contact with surfaces (36, 38). We confirmed that the root colonization phenotypes of Δ*gacS*, Δ*gacA*, and Δ*fliF* were complemented by ectopic expression of arabinose-inducible forms of each mutated gene (Fig. S2B). We then used epistasis analysis to determine whether the *gac* and flagellar genes contributed to a unified pathway controlling root attachment. A Δ*fliF* Δ*gacA* double mutant displayed a root colonization defect comparable to the Δ*gacA* single mutant, indicating that the *gac* system is required for enhanced root attachment in flagellar mutants (Fig. 4A, Fig. S3). To determine whether differences in root attachment correlate with altered c-di-GMP levels, which are known to promote bacterial adhesion during responses to surface contact, we quantified c-di-GMP levels using a fluorescent reporter (39). The Δ*fliF* mutant showed elevated c-di-GMP and the Δ*gacA* mutant exhibited lower c-di-GMP concentrations than wild-type. The Δ*fliF* Δ*gacA* double mutant displayed c-di-GMP levels matching the wild-type phenotype and confirming that the hyperadhesive, high-c-di-GMP state of Δ*fliF* depends on *gacA*.

**Figure 4.**
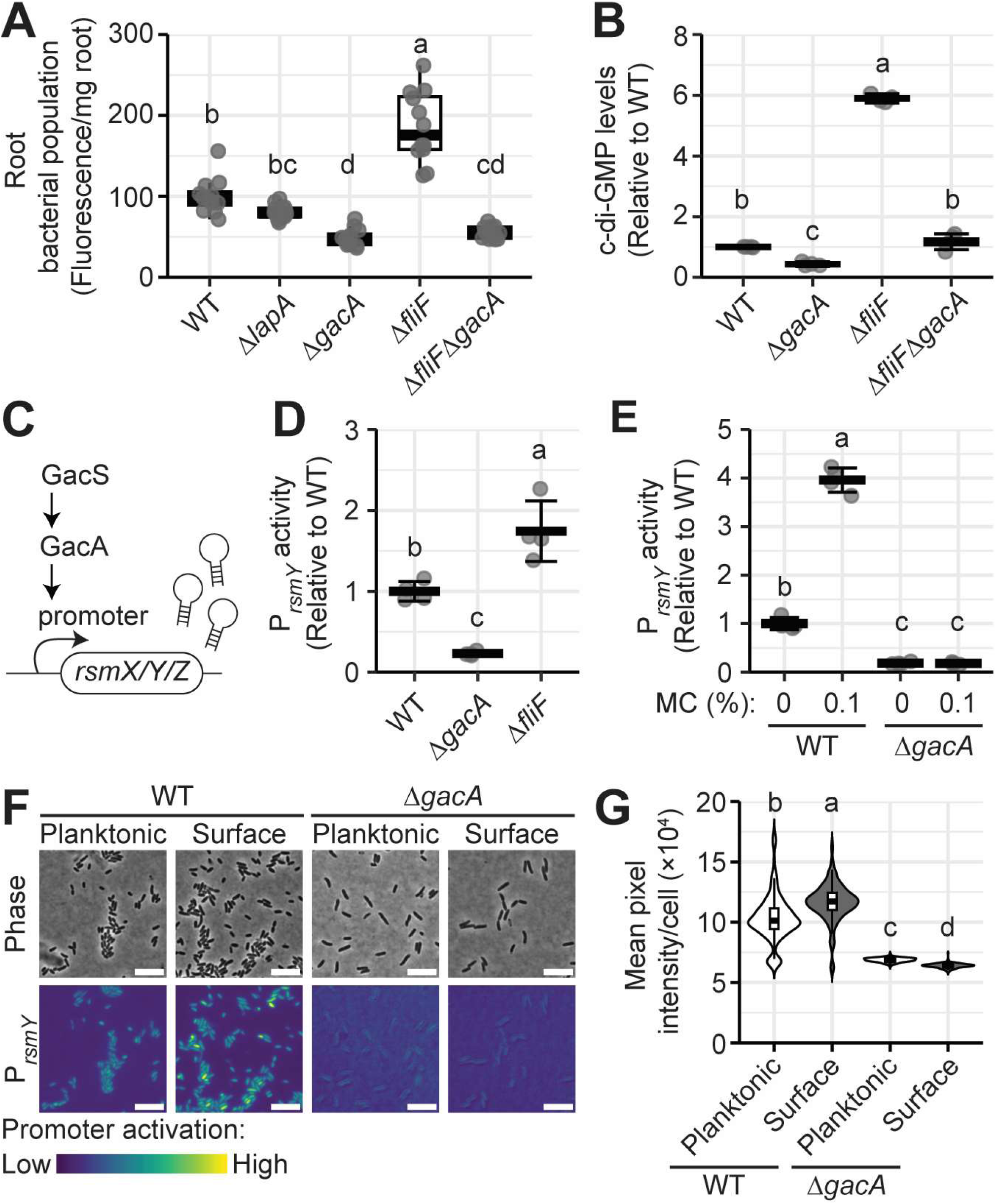
Flagellar-dependent surface sensing activates the *gac* system (A) Root colonization efficiencies of key mutants. Top and bottom sides of the boxes represent the third and first quartiles of the value distribution, respectively. Lines extending from the boxes denote the extreme values within 1.5 times the interquartile range. Data are pooled from three independent replicates. (B) Intracellular c-di-GMP levels of strains from panel A. Reporter values were normalized to wild type. (C) Overview of the *gac* signaling system in *Pseudomonads.* GacS activates the response regulator GacA, which in turn directly induces expression of the small RNAs *rsmX*, *rsmY*, and *rsmZ*. In D-G, the effect of surface sensing on *gac* activity was evaluated via P*_rsmY_* expression by increasing medium viscosity with methyl cellulose (MC) (D), disrupting flagellar assembly (Δ*fliF*) (E), or comparing planktonic and surface-attached cells (F and G). In F, scale bars represent 5 μm. In B, D, and E, a representative replicate of three individual experiments is shown. Data are means ± SD. In all panels letters represent statistical significance determined by one-way ANOVA and Tukey’s posthoc test (*P* < 0.05).

Mutations that disrupt flagellar assembly enhance biofilm formation in several bacteria by activating surface sensing pathways (36, 38). We predicted that the Δ*fliF* mutation enhances adhesion by mimicking surface contact, which in turn activates the *gac* system. We monitored transcriptional reporters for the three small RNAs that are directly activated by GacA∼P in Pf-5 (*rsmX*, *rsmY*, and *rsmZ*) under conditions designed to induce surface sensing (Fig. 4C) (40, 41). Surface sensing was stimulated by increasing medium viscosity with methyl cellulose (42), by disrupting flagellar assembly using the Δ*fliF* mutation, or by incubating cells on an agar surface. *gac* activity was elevated across all conditions where surface sensing was stimulated (Fig. 3D-G). Similar trends were observed for all three *rsm* reporters (Fig. S4). Together, these findings support a model in which physical contact with surfaces causes the flagellum to activate the *gac* system.

### The *gac* system integrates physical and chemical cues to control surface attachment

To evaluate how the attachment phenotypes we measured on root tissue compare to abiotic surface attachment, we quantified biofilm formation in microtiter plates using a crystal violet (CV) staining assay. We observed a different attachment pattern on an abiotic surface than we measured on root tissue. The Δ*fliF* mutant remained hyperadhesive on abiotic surfaces, and a Δ*fliF* Δ*gacA* double mutant still lost the enhanced adhesion of Δ*fliF* alone (Fig. 5A). However, the Δ*gacA* single mutant and the Δ*fliF* Δ*gacA* double mutant showed identical levels of CV staining to the wild-type strain. These results indicate the *gac* system plays a stronger role in attachment to tissue than to abiotic surfaces.

**Figure 5.**
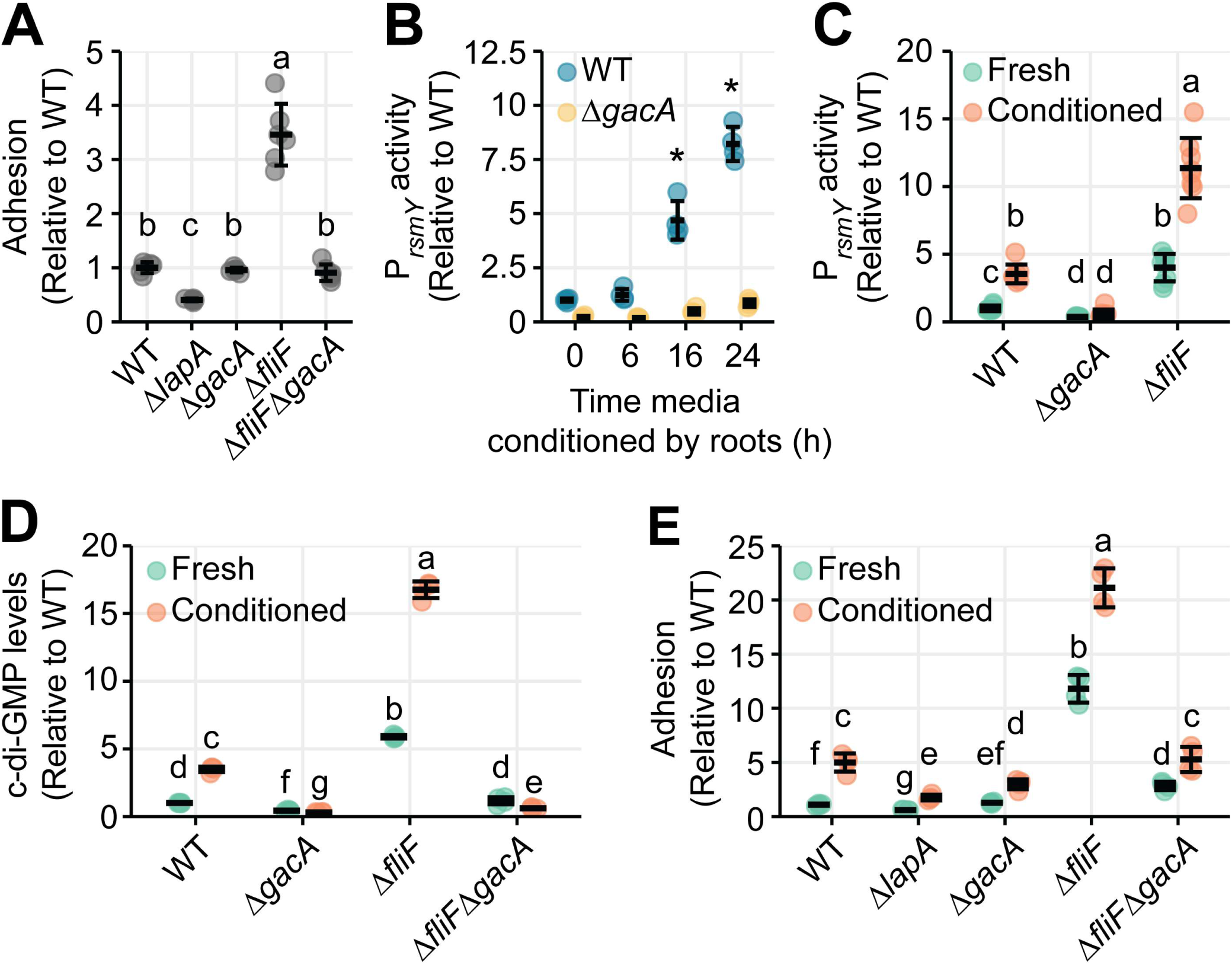
The *gac* system integrates physical and chemical cues to control surface attachment (A) Crystal violet (CV)-based adhesion assay evaluating attachment to an abiotic surface. (B) *gac* system activity quantified using a fluorescent reporter for *rsmY* evaluating the effect of conditioning time of media by roots on *gac* system activity. (C–E) Integration of chemical and mechanical inputs by the *gac* system. Indicated strains were analyzed under conditions providing mechanical input alone (Δ*fliF*), chemical input alone (root-conditioned media), or both inputs combined. (C) *gac* system activity quantified via the *rsmY* fluorescent reporter. (E) Intracellular c-di-GMP levels. (F) CV-based adhesion assay. *gac* system activity levels, c-di-GMP levels, and adhesion were normalized to wild-type in unconditioned (Fresh) media. A representative replicate of three independent experiments is shown. Data represent means ± SD. In all panels, different letters indicate statistically significant differences determined by one-way ANOVA with Tukey’s post hoc test (*P* < 0.05). In panel B, asterisks indicate statistically significant differences determined by pairwise *t*-tests comparing each time point to the preceding time point within the same strain.

We predicted that differences in colonization efficiencies of *gac* mutants on root tissues compared to abiotic surfaces could be explained by the presence of exudates that accumulate in tissue culture medium (43, 44). We measured *gac* reporter activity in media that had been conditioned by roots for 6, 16, and 24 hours. Activation of all three *gac* reporters increased with longer root conditioning times. No activation was observed in the Δ*gacA* mutant (Fig. 5B, Fig. S5A and C). These results demonstrate that root-secreted molecules stimulate *gac* signaling.

We next tested whether the chemical signals from root exudates and physical signals from the flagellum activate the *gac* system in an additive manner or through a sequential logic in which one cue is required before the other can be detected. To distinguish these possibilities, we examined *gac* activity, c-di-GMP levels, and surface attachment in a CV assay, under conditions with surface sensing activated or chemical cues alone, or both inputs together. When we combined the two signals using root-conditioned media and a Δ*fliF* mutant, we observed higher levels of *gac* reporter activation, higher c-di-GMP levels, and increased adhesion, compared to conditions with either input alone (Fig. 5C-E, Fig. S5B and D). This enhancement indicates that the *gac* system integrates physical and chemical signals to promote root colonization.

### Activating flagellar surface sensing enhances competitive root colonization

Rhizosphere colonization efficiency is thought to be driven by competitive interactions among soil organisms. We predicted that activating surface responses and *gac* signaling using the Δ*fliF* mutation would provide Pf-5 with a competitive advantage over other bacteria during root attachment. We performed a root attachment assay in the presence of THOR, a synthetic bacterial community composed of three members isolated from the soybean rhizosphere (45).

Co-inoculation with THOR reduced the attachment efficiency of wild-type Pf-5 by ∼40%, while the Δ*gacA* mutant remained at low abundance (Fig. 6A). In contrast, the Δ*fliF* mutant maintained elevated attachment efficiency in the presence of THOR. These results show that enhanced activation of attachment programs in the Δ*fliF* mutant is sufficient to overcome bacterial competition in the root environment.

**Figure 6.**
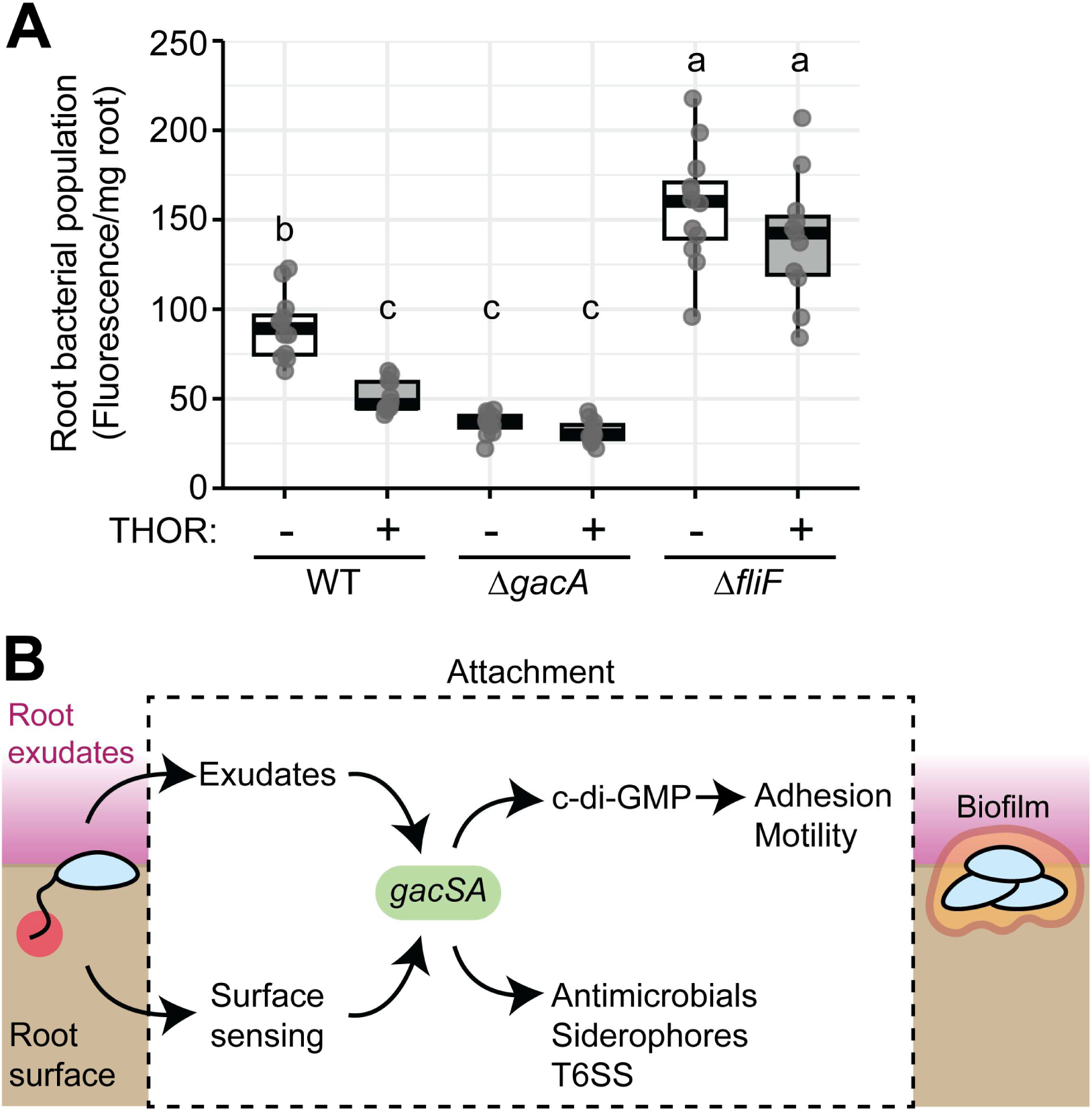
Activating flagellar surface sensing enhances competitive root colonization (A) Root colonization phenotypes in the presence of a synthetic bacterial community. Root colonization was quantified by recovering the root-attached fraction of cells and measuring its emitted fluorescence. Bold lines within each box represent median values. Top and bottom sides of the boxes represent the third and first quartiles of the value distribution, respectively. Lines extending from the boxes denote the extreme values within 1.5 times the interquartile range. Data are pooled from three independent replicates. (B) The *gac* system additively integrates physical and chemical cues to coordinately enhance adhesion and reduce motility through c-di-GMP, while simultaneously activating competitive traits, together enabling effective root attachment.

## Discussion

Previous studies have used whole plants to identify bacterial genes that promote persistent colonization of roots (7). These efforts have yielded foundational insights into plant-microbe interactions in the rhizosphere, but the crucial stage of early root attachment has remained less characterized. In this study, we developed an *in vitro* root tissue culture system to specifically target the early stages of bacterial root attachment. Propagating large quantities of roots in liquid medium eliminates the time and logistical effort needed for whole-plant experiments. The higher throughput and precise experimental control of a tissue culture system allowed us to identify phenotypes that were not apparent in previous studies (7). For example, only a subset of the attachment determinants we found were identified when Pf-5 mutants were inoculated onto wheat and cotton seeds, grown in a sand/perlite mix, and recovered after one week (21). More broadly, results of many long-duration root colonization Tn-Seq studies are dominated by genes involved in metabolism, transport, cell wall biogenesis, and stress adaptation rather than classical biofilm factors (7, 14, 16, 18, 20–23, 25, 46, 47). These contrasts highlight the complex temporal progression of root attachment. Individual genes likely play stage-dependent roles, and early advantages may later become neutral or even detrimental as colonization progresses. Targeting the early attachment phase specifically led to the identification of new signaling connections that influence attachment to roots.

The *gacSA* two component signaling system emerged as a crucial determinant of root attachment in our study. While the *gac* system is known to influence a large set of behaviors in *Pseudomonads,* there are conflicting reports about its role in biofilm formation and tissue colonization. *gac* mutants generally show reduced adhesion to abiotic surfaces in crystal violet assays (48–56). However, other studies indicate that abiotic surface adhesion phenotypes of *gac* mutants are inconsistent and may depend on specific culture conditions (57–59; This study). In root colonization assays with whole plants, the consequences of *gac* disruption are strongly shaped by competition. *gac* mutants often do not show impaired root colonization and may even outperform wild type in competition (60–62), reflecting a public goods tradeoff (55, 63). In contrast, *gac* mutants show a competitive disadvantage in natural soil or mixed-community environments (64–66). While previous Tn-Seq colonization studies that selected for competitive growth identified *gac*, it was not among the strongest contributors to root colonization (31, 46). In our study, *gacS* and *gacA* mutants had by far the strongest adhesion defects of any gene in our early root attachment system. We propose that *gac* activation is crucial for promoting tissue adhesion specifically during the earliest stages of root contact.

Our study also illuminates a novel function for flagella in the root colonization sequence. Flagella have long been recognized as surface colonization factors, but previous root attachment studies have identified a range of phenotypes for flagellar mutants. Root attachment defects, neutral colonization phenotypes and hyper-colonization of roots have all been reported (16, 47, 64, 67, 68). Variations likely arise because specific experimental methods inadvertently emphasize different stages of root colonization, but previous studies have mostly interpreted flagellar phenotypes through the lens of motility. Colonization defects are attributed to an inability to migrate efficiently to the root surface, and hyper-colonization is assumed to arise when cells become “trapped” on the root because they cannot swim away. By focusing our tissue colonization system specifically on the early attachment phase, we provide evidence for flagella functioning as surface sensors. The enhanced adhesion and elevated c-di-GMP levels of the Δ*fliF* mutant are consistent with studies showing that disrupting flagellar assembly mimics surface contact, and we could recapitulate the effects of the Δ*fliF* mutant in wild-type cells by physically obstructing the flagellum (Fig. 4D-G, Fig. S4). Thus, our results provide evidence that surface sensing mechanisms identified for abiotic surfaces are preserved in interactions with living tissue.

Root exudates emerged as a key differentiator between abiotic surfaces and root tissue. Root conditioned medium stimulated *gac* signaling, c-di-GMP production, and adhesion in a *gac-*dependent manner. While we do not yet know the exact exudate compound(s) responsible for this effect, our findings underscore how *gac* phenotypes seem to be particularly sensitive to chemical changes. Perhaps tuning the system to subtle chemical changes potentiates tissue colonization. More broadly, exudates are by large viewed as chemoattractants that promote motility toward the root (12). Our data supports an expanded view whereby exudates also function as local signals that directly enhance root attachment (13, 69). This finding underscores a fundamental difference between abiotic and tissue surfaces. Living tissues modify the chemical environment through secretion in a way that cannot be reproduced on abiotic substrates, and we propose that the *gac* system among *Pseudomonas fluorescens* species complex strains has evolved to incorporate chemical cues from roots into the biofilm activation program.

Two-component and phosphorelay systems frequently operate as signal-integration nodes by enforcing coincident detection of multiple signals before committing to irreversible responses (70–72). Our data place *gacSA* within this framework as an integrator of both chemical and physical cues from roots. While many studies have examined how the sensor kinase GacS is activated, there is still no consensus on the activating ligand(s) (73). Our data show that both surface contact sensing through the flagella and chemical cues in exudates activate the *gac* system on roots, but we do not yet know if these are direct activators of GacS.

GacS activation triggers *gacA*-dependent expression of *rms* sRNAs that go on to affect genes for biofilm formation, plant growth promotion, siderophore production and antimicrobial production (37, 74, 75). Our data indicates that root attachment requires a *gac*-depedent increase in c-di-GMP levels, and we indeed identified two c-di-GMP metabolic genes that promote root attachment in our screen (*PFL_4532*, *PFL_5779*; Fig. 3). c-di-GMP binds transcriptional regulators that shift cells to a sessile state by repressing motility while activating adhesins (76–78). The absence of strong hits in specific adhesins in our screen likely reflects redundancy among attachment factors and suggests that regulation occurs primarily at the level of c-di-GMP signaling rather than direct transcriptional activation by *rsm* regulators. In many bacteria, surface colonization follows a relatively simple pathway in which surface contact elevates cyclic-di-GMP and activates adhesins and biofilm matrix components (79, 80). In contrast, in the *Pseudomonas fluorescens* species complex this transition is embedded within the global *gac* regulatory system. Our results suggest that this expanded regulation reflects the complexity of the rhizosphere, where root colonization likely requires coordinated activation of adhesion together with traits involved in competition and plant interaction (Fig. 6B). An important question is whether *gac* activation engages these programs proportionally or can bias them toward distinct outputs. One possibility is that differences in the magnitude or duration of *gac* signaling are encoded through the kinetics of the *rsm* sRNAs, allowing the *gac* system to tune the balance between adhesion and competitive behaviors depending on environmental conditions during root colonization.

Treatments with beneficial rhizobacteria perform poorly under field conditions in part because competition from native microbiota limits effective root colonization (81, 82). This ecological constraint means that even large inoculum doses or recurrent applications often provide limited long-term enhancement of plant performance. Current efforts to enhance colonization focus largely on improving delivery through strategies that position bacteria near emerging roots and by isolating new strains that may be better suited for competing in the rhizosphere. While these approaches can improve early establishment, they do not directly address how bacteria transition from bulk soil to the root surface. By showing that flagellar surface sensing stimulates the *gac* signaling system, our work suggests a new route to design more persistent bioinoculants. Rather than relying on genetic engineering or reformulating products, it could be possible to select for naturally occurring variants with reduced flagellar synthesis under the hypothesis that reduced motility will enhance root attachment at high inoculation titers. Combining such surface sensing-optimized strains with existing delivery technologies like seed coatings could yield inoculants that deliver more consistent yield enhancement in the field.

## Materials and Methods

### Growth conditions and genetic manipulations

Growth media were supplemented with 1.5% (w/v) agar, 300 µM diaminopimelic acid (DAP), 50 µg/mL gentamycin, 100 µg/mL carbenicillin, and 50 µg/mL kanamycin when necessary. *E*. *coli* was cultured in LB medium at 37°C, *P*. *protegens* in King’s B medium (KBM) at 30°C, *A*. *rhizogenes* in Nutrient broth at 30°C. Plasmids were introduced into *P*. *protegens* Pf-5 by biparental conjugation with WM3064 as the donor strain. Gene deletions or insertions were generated using a two-step approach with a *sacB-*based counterselection. Mutants were complemented by genomic integration of the deleted gene under the arabinose promoter using the Tn*7* transposon (83). Mutants were fluorescently-tagged by inserting an mVenus expression cassette into the CTX *att* site or *glmS* site (83, 84). Plasmids and strains used in this study are listed in Table S2 and Table S3, respectively.

### Establishment of tomato hairy root cultures

Tomato (*Solanum lycopersicum* cv. Rio-Grande) seedlings were transformed with *Agrobacterium rhizogenes* 15834 to generate hairy roots as previously described (85). To generate a root biomass, the most vigorous roots, identified by their bright white color, high density of root hairs, and length, were excised and transferred to 250 mL flasks containing 50 mL of 0.5× MS medium (pH 5.8) with 3% sucrose and supplemented with 200 µg/mL carbenicillin and 200 µg/mL cefotaxime to suppress *Agrobacterium* growth. Flasks were incubated in the dark at 25 °C with shaking at 75 rpm, and the medium was refreshed weekly. After 2-3 weeks, when roots had filled the entire flask volume, they were cut and transferred into microplate wells for attachment assays.

### Tn-*Himar* mutant library construction and mapping

Construction and mapping of the Pf-5 transposon library was performed based on the method developed by Wetmore *et al*. (34). Mid-log phase cells from cultures of the APA_752 barcoded transposon pool grown in LB medium supplemented with kanamycin and DAP and mid-log phase Pf-5 cells from cultures grown in KBM were collected by centrifugation, washed, mixed in a 1:1 ratio, and spotted on a KBM agar plate containing DAP. After incubation for 24 h at 30°C, cells were scraped from the plate, resuspended in KBM medium, and spread onto 20 150-mm KBM plates containing kanamycin, followed by incubation at 30°C for 3 days. Colonies were scraped into KBM medium and used to inoculate a KBM culture containing 25 μg/mL kanamycin. The culture was grown for three doublings, glycerol was added to 15%, and 1 mL aliquots were frozen at-80°C. Mapping genomic insertion positions was performed as described previously (86). The features of the obtained library are: 57,304,857 total reads, 25,081,887 mapped reads in pool, 784,810 mapped barcodes in pool, 161,378 distinct insertion sites, 5,816 coding genes with central insertions, 53 median strains per hit protein, 81.7 mean strains per hit protein.

### Adhesion profiling of barcoded Tn-*Himar* mutant libraries

Adhesion profiling was performed based on the method developed by Hershey *et al*. (86). Flasks containing hairy roots were prepared by refreshing the medium 48 h prior to inoculation, allowing root-secreted compounds to accumulate. A 750 μL inoculum of the barcoded transposon library was added to a 250 mL flask containing 75 mL of medium and hairy roots, and flasks were incubated for 24 h at 25°C with shaking at 75 rpm to allow bacterial attachment. Following incubation, 750 μL of the culture was transferred into a new flask with hairy roots, while cells from an additional 30 mL of culture were collected by centrifugation and stored at −20°C for BarSeq analysis. This procedure was performed five times for a total of five sequential passaging rounds. Parallel cultures were grown in flasks containing root-conditioned medium but no hairy roots, serving as a reference condition. Genomic DNA was extracted from cell pellets and served as templates for amplifying the barcodes in each sample using indexed primers (34). Amplicons were purified using AMPure XP magnetic beads (Beckman Coulter) and pooled for multiplexed sequencing. Next, 50-bp single-end reads were collected on an Illumina NovaSeq X Plus sequencer. MultiCodes.pl, combineBarSeq.pl, and BarSeqR.pl were used to determine fitness by comparing the log_2_ ratios of barcode counts in each sample over the counts from nonselective growth without hairy roots. For each mutant, the pracma R package was used to calculate the area under the curve of fitness scores across passages to rank mutants according to their hypo-or hyper-adhesive phenotypes.

### Crystal violet staining assay

Overnight cultures were resuspended in 0.5× MS medium (pH 5.8) supplemented with 3% sucrose and 20 mM succinate, which was either freshly prepared or conditioned with hairy roots for 24 h. Resuspensions were normalized to OD_600_ 0.002, and 450 μL was added to each well of a 48-well microtiter plate. Plates were incubated shaking for 24 h at 30°C and 200 rpm. OD_600_ was measured to account for growth differences. Culture broth was discarded and the wells were washed by dipping the microtiter plate in a bin with tap water. 500 μL of 0.1% (w/v) crystal violet solution was added to each well, and the plate was shaken for 30 min before the dye was discarded and wells washed in the same manner. The retained dye was dissolved in 500 μL of absolute ethanol by shaking for 10 min. OD_575_ was measured in each well and normalized to OD_600_.

### Measurement of c-di-GMP

Strains were transformed with the pConRef-2H12.D11 c-di-GMP reporter plasmid (39). Overnight cultures were resuspended in 0.5× MS medium (pH 5.8) supplemented with 3% sucrose and 20 mM succinate. MS media was either freshly prepared or conditioned with hairy roots for 24 h. Cell suspensions were normalized to OD_600_ 0.02, and 200 μL was dispensed into wells of a 96-well microtiter plate. Plates were incubated shaking for 24 h at 30 °C and 200 rpm, after which fluorescence was measured using a BioTek Synergy H1 microplate reader for cdGreen2.1 (Excitation: 470 nm, emission: 515 nm) and normalized to the reference mScarlet-I (Excitation: 575 nm, emission: 632 nm).

### Measurement of *gac* activity reporters

Strains were grown overnight in 0.5× MS medium (pH 5.8) supplemented with 3% sucrose and 20 mM succinate. To evaluate the effects of the Δ*fliF* mutation and methyl cellulose on reporter activity, overnight cultures were diluted 100-fold into 96-well plates containing 200 μL of the indicated medium per well and incubated for 24 h. Plates were shaken when assessing the effect of the Δ*fliF* mutation and incubated statically for methyl cellulose. Fluorescence was measured using a BioTek Synergy H1 microplate reader (mVenus; Excitation: 500 nm; emission: 540 nm) and normalized to OD_600_. For microscopy, overnight cultures were diluted 10-fold, and 2 µL of cells were placed on a 2% agarose pad to immobilize the cells. Cells were imaged immediately to quantify reporter activity in planktonic cells and imaged again after 1 h to measure reporter activity in surface-sensing cells. Microscopy was performed using a Nikon Ti-E inverted microscope equipped with an Orca Fusion BT digital CMOS camera (Hamamatsu). Fluorescence images were collected using a Prior Lumen 200 metal halide light source and a YFP-specific filter set (Chroma) with identical exposure times and laser intensity. Image analysis was performed with the Fiji software.

### Root attachment assays

Hairy roots were excised into segments and placed in wells of a 48-well plate with 0.5× MS medium (pH 5.8) supplemented with 3% sucrose and 20 mM succinate conditioned by roots for 24h. Wells were inoculated with the indicated strains at OD_600_ 0.02. Plates were incubated at 24°C with shaking at 75 rpm. To qualitatively assess bacterial root attachment, roots were washed three times within the plate wells and imaged for fluorescence using the Invitrogen™ iBright™ CL1500 Imaging System (Excitation: 455-485 nm, emission: 508-557 nm). To quantitatively assess root attachment, roots were transferred to microcentrifuge tubes containing 1 mL of 0.5× MS medium. Root-associated bacteria were detached by vortexing for 1 min, and the resulting suspensions were serially diluted and plated on LB agar for CFU enumeration, or had their fluorescence measured using a BioTek Synergy H1 microplate reader (mVenus; Excitation: 500 nm, emission: 540 nm).

### Bacterial growth curve measurement

Overnight cultures were grown overnight in LB medium at 30 °C and adjusted to OD_600_ 0.02 in indicated media. 200 µL aliquots were dispensed into 96-well plates, with five technical replicates per strain. Plates were incubated at 30 °C with shaking in a BioTek Synergy H1 microplate reader, and OD_600_ was recorded at 30-min intervals for the duration of the experiment.

## Data availability

The sequence data used to map the barcoded Tn-*Himar* library and the barcoded amplicon sequences collected after passaging in plant roots have been deposited in the NCBI Sequence Read Archive (SRA) under the PRJNA1440512 project accession number. Strains, plasmids, and details of their construction are available upon request.

## Acknowledgements

This work was supported by a National Institutes of Health award R35GM150652 to D.M.H, Beckman Young Investigator award to D.M.H., and startup funds from the University of Wisconsin – Madison to D.M.H. This research was supported by BARD, the United States - Israel Binational Agricultural Research and Development Fund, Vaadia-BARD Postdoctoral Fellowship Award No. FI-638-2024. We thank Or Sharon for technical assistance with preparing and uploading data to the NCBI SRA.

## References

1. Sauer K, Stoodley P, Goeres DM, Hall-Stoodley L, Burmølle M, Stewart PS, Bjarnsholt T. 2022. The biofilm life cycle: expanding the conceptual model of biofilm formation. Nat Rev Microbiol 20:608–620.

2. Flemming H-C, Wuertz S. 2019. Bacteria and archaea on Earth and their abundance in biofilms. Nat Rev Microbiol 17:247–260.

3. Baxter KJ, Sas E, Clark KB, Walsh M, Pradeep N, Batool A, Naney C, Vargas Cruz MA, Kennerdale N, Das K, Shi Z, Kelam A, Verma V, Simões MF, Neefs D, Ravichandran V, Tirumalai MR, Barcenilla BB, Macori G, Gonzalez E, Sikes B, Karouia F, Brereton NJB. 2026. Biofilms: from the cradle of life to life support. npj Biofilms Microbiomes 12:11.

4. Grari O, Ezrari S, El Yandouzi I, Benaissa E, Ben Lahlou Y, Lahmer M, Saddari A, Elouennass M, Maleb A. 2025. A comprehensive review on biofilm-associated infections: Mechanisms, diagnostic challenges, and innovative therapeutic strategies. The Microbe 8:100436.

5. Liu Y, Shi A, Chen Y, Xu Z, Liu Y, Yao Y, Wang Y, Jia B. 2025. Beneficial microorganisms: Regulating growth and defense for plant welfare. Plant Biotechnology Journal 23:986–998.

6. Rumbaugh KP, Whiteley M. 2025. Towards improved biofilm models. Nat Rev Microbiol 23:57–66.

7. Torres M, Paszti S, Eberl L. 2024. Shedding light on bacteria–host interactions with the aid of TnSeq approaches. mBio 15:e00390–24.

8. Pearce D, Bazin MJ, Lynch JM. 1995. The rhizosphere as a biofilm, p. 207–220. Cambridge University Press, Cambridge.

9. Liu Y, Xu Z, Chen L, Xun W, Shu X, Chen Y, Sun X, Wang Z, Ren Y, Shen Q, Zhang R. 2024. Root colonization by beneficial rhizobacteria. FEMS Microbiol Rev 48:fuad066.

10. Bhattacharyya A, Mavrodi O, Bhowmik N, Weller D, Thomashow L, Mavrodi D. 2023. Bacterial biofilms as an essential component of rhizosphere plant-microbe interactions. Methods Microbiol 53:3–48.

11. Robert CAM, Himmighofen P, McLaughlin S, Cofer TM, Khan SA, Siffert A, Sasse J. 2025. Environmental and biological drivers of root exudation. Annual Review of Plant Biology 76:317–339.

12. Yang C-X, Chen S-J, Hong X-Y, Wang L-Z, Wu H-M, Tang Y-Y, Gao Y-Y, Hao G-F. 2025. Plant exudates-driven microbiome recruitment and assembly facilitates plant health management. FEMS Microbiol Rev 49:fuaf008.

13. Bhattacharyya A, Mavrodi O, Bhowmik N, Weller D, Thomashow L, Mavrodi D. 2023. Bacterial biofilms as an essential component of rhizosphere plant-microbe interactions. Methods Microbiol 53:3–48.

14. Liu Z, Beskrovnaya P, Melnyk RA, Hossain SS, Khorasani S, O’Sullivan LR, Wiesmann CL, Bush J, Richard JD, Haney CH. 2018. A genome-wide screen identifies genes in rhizosphere-associated *Pseudomonas* required to evade plant defenses. mBio 9:10.1128/mbio.00433-18.

15. Torres M, Jiquel A, Jeanne E, Naquin D, Dessaux Y, Faure D. 2022. *Agrobacterium tumefaciens* fitness genes involved in the colonization of plant tumors and roots. New Phytologist 233:905–918.

16. do Amaral FP, Tuleski TR, Pankievicz VCS, Melnyk RA, Arkin AP, Griffitts J, Tadra-Sfeir MZ, Maltempi de Souza E, Deutschbauer A, Monteiro RA, Stacey G. 2020. Diverse bacterial genes modulate plant root association by beneficial bacteria. mBio 11:10.1128/mbio.03078-20.

17. Cole BJ, Feltcher ME, Waters RJ, Wetmore KM, Mucyn TS, Ryan EM, Wang G, Ul-Hasan S, McDonald M, Yoshikuni Y, Malmstrom RR, Deutschbauer AM, Dangl JL, Visel A. 2017. Genome-wide identification of bacterial plant colonization genes. PLOS Biology 15:e2002860.

18. Knights HE, Ramachandran VK, Jorrin B, Ledermann R, Parsons JD, Aroney STN, Poole PS. 2024. *Rhizobium* determinants of rhizosphere persistence and root colonization. ISME J 18:wrae072.

19. Torres M, Price MN, Khasanova A, Kosina SM, Zhalnina K, Northen TR, Deutschbauer AM. 2025. Bacterial fitness for plant colonization is influenced by plant growth substrate. New Phytologist 248:3168–3190.

20. Sivakumar R, Ranjani J, Vishnu US, Jayashree S, Lozano GL, Miles J, Broderick NA, Guan C, Gunasekaran P, Handelsman J, Rajendhran J. 2019. Evaluation of INSeq to identify genes essential for *Pseudomonas aeruginosa* PGPR2 corn root colonization. G3 Genes|Genomes|Genetics 9:651–661.

21. Ghaly TM, Fabian BK, Vick SHW, Foster C, Asher AJ, Hassan KA, Elbourne LDH, Paulsen IT, Tetu SG. 2025. Genetic drivers of plant root colonisation by the biocontrol agent *Pseudomonas protegens* Pf-5. Environmental Microbiology Reports 17:e70179.

22. Cheng X, Etalo DW, van de Mortel JE, Dekkers E, Nguyen L, Medema MH, Raaijmakers JM. 2017. Genome-wide analysis of bacterial determinants of plant growth promotion and induced systemic resistance by *Pseudomonas fluorescens*. Environmental Microbiology 19:4638–4656.

23. Pranav PS, Sivakumar R, Suvekbala V, Rajendhran J. 2024. Genome-wide identification of root colonization fitness genes in plant growth promoting *Pseudomonas asiatica* employing transposon-insertion sequencing. Ann Microbiol 74:40.

24. Taylor TB, Silby MW, Jackson RW. 2025. Pseudomonas fluorescens. Trends in Microbiology 33:250–251.

25. Zboralski A, Filion M. 2020. Genetic factors involved in rhizosphere colonization by phytobeneficial *Pseudomonas* spp. Computational and Structural Biotechnology Journal 18:3539–3554.

26. Krell T, Matilla MA. 2024. Pseudomonas aeruginosa. Trends in Microbiology 32:216–218.

27. Collins AJ, Smith TJ, Sondermann H, O’Toole GA. 2020. From input to output: The Lap/c-di-GMP biofilm regulatory circuit. Annual Review of Microbiology 74:607–631.

28. Loper JE, Kobayashi DY, Paulsen IT. 2007. The genomic sequence of *Pseudomonas fluorescens* Pf-5: Insights into biological control. Phytopathology® 97:233–238.

29. Paulsen IT, Press CM, Ravel J, Kobayashi DY, Myers GSA, Mavrodi DV, DeBoy RT, Seshadri R, Ren Q, Madupu R, Dodson RJ, Durkin AS, Brinkac LM, Daugherty SC, Sullivan SA, Rosovitz MJ, Gwinn ML, Zhou L, Schneider DJ, Cartinhour SW, Nelson WC, Weidman J, Watkins K, Tran K, Khouri H, Pierson EA, Pierson LS, Thomashow LS, Loper JE. 2005. Complete genome sequence of the plant commensal *Pseudomonas fluorescens* Pf-5. Nat Biotechnol 23:873–878.

30. Jing X, Cui Q, Li X, Yin J, Ravichandran V, Pan D, Fu J, Tu Q, Wang H, Bian X, Zhang Y. 2020. Engineering *Pseudomonas protegens* Pf-5 to improve its antifungal activity and nitrogen fixation. Microbial Biotechnology 13:118–133.

31. Ghaly TM, Fabian BK, Vick SHW, Foster C, Asher AJ, Hassan KA, Elbourne LDH, Paulsen IT, Tetu SG. 2025. Genetic drivers of plant root colonisation by the biocontrol agent *Pseudomonas protegens* Pf-5. Environmental Microbiology Reports 17:e70179.

32. Zhao Q, Wang R, Song Y, Lu J, Zhou B, Song F, Zhang L, Huang Q, Gong J, Lei J, Dong S, Gu Q, Borriss R, Gao X, Wu H. 2024. Pyoluteorin-deficient *Pseudomonas protegens* improves cooperation with *Bacillus velezensis*, biofilm formation, co-colonizing, and reshapes rhizosphere microbiome. npj Biofilms Microbiomes 10:145.

33. Kamilova F, Kravchenko LV, Shaposhnikov AI, Makarova N, Lugtenberg B. 2006. Effects of the tomato pathogen *Fusarium oxysporum* f. sp. *radicis-lycopersici* and of the biocontrol bacterium *Pseudomonas fluorescens* WCS365 on the composition of organic acids and sugars in tomato root exudate. MPMI 19:1121–1126.

34. Wetmore KM, Price MN, Waters RJ, Lamson JS, He J, Hoover CA, Blow MJ, Bristow J, Butland G, Arkin AP, Deutschbauer A. 2015. Rapid quantification of mutant fitness in diverse bacteria by sequencing randomly bar-coded transposons. mBio 6:e00306–15–15.

35. Niehaus TD, Elbadawi-Sidhu M, Crécy-Lagard V de, Fiehn O, Hanson AD. 2017. Discovery of a widespread prokaryotic 5-oxoprolinase that was hiding in plain sight. Journal of Biological Chemistry 292:16360–16367.

36. Laventie B-J, Jenal U. 2020. Surface Sensing and adaptation in bacteria. Annual Review of Microbiology 74:735–760.

37. Song H, Li Y, Wang Y. 2023. Two-component system GacS/GacA, a global response regulator of bacterial physiological behaviors. Engineering Microbiology 3:100051.

38. Dufrêne YF, Persat A. 2020. Mechanomicrobiology: how bacteria sense and respond to forces. Nat Rev Microbiol 18:227–240.

39. Kaczmarczyk A, Van Vliet S, Jakob RP, Teixeira RD, Scheidat I, Reinders A, Klotz A, Maier T, Jenal U. 2024. A genetically encoded biosensor to monitor dynamic changes of c-di-GMP with high temporal resolution. Nat Commun 15:3920.

40. Kay E, Dubuis C, Haas D. 2005. Three small RNAs jointly ensure secondary metabolism and biocontrol in *Pseudomonas fluorescens* CHA0. Proceedings of the National Academy of Sciences 102:17136–17141.

41. Zha D, Xu L, Zhang H, Yan Y. 2014. The two-component GacS-GacA system activates lipA translation by RsmE but Not RsmA in *Pseudomonas protegens* Pf-5. Applied and Environmental Microbiology 80:6627–6637.

42. Pijper A. 1947. Methylcellulose and bacterial motility. J Bacteriol 53:257–269.

43. Joller C, Waelchli J, Schlaepfer J, Schlaeppi K. 2026. Paralleled dynamics of *Arabidopsis* root exudation and syncom assembly in a controlled environment. bioRxiv 10.64898/2026.01.29.702624.

44. McLaughlin S, Zhalnina K, Kosina S, Northen TR, Sasse J. 2023. The core metabolome and root exudation dynamics of three phylogenetically distinct plant species. Nat Commun 14:1649.

45. Lozano GL, Bravo JI, Garavito Diago MF, Park HB, Hurley A, Peterson SB, Stabb EV, Crawford JM, Broderick NA, Handelsman J. 2019. Introducing THOR, a model microbiome for genetic dissection of community behavior. mBio 10: pp.10–1128.

46. Cole BJ, Feltcher ME, Waters RJ, Wetmore KM, Mucyn TS, Ryan EM, Wang G, Ul-Hasan S, McDonald M, Yoshikuni Y, Malmstrom RR, Deutschbauer AM, Dangl JL, Visel A. 2017. Genome-wide identification of bacterial plant colonization genes. PLOS Biology 15:e2002860.

47. Wallner A, Busset N, Lachat J, Guigard L, King E, Rimbault I, Mergaert P, Béna G, Moulin L. 2022. Differential genetic strategies of *Burkholderia vietnamiensis* and *Paraburkholderia kururiensis* for root colonization of *Oryza sativa* subsp. *japonica* and *O. sativa* subsp. *indica*, as revealed by transposon mutagenesis sequencing. Applied and Environmental Microbiology 88:e00642–22.

48. López-Sánchez A, Leal-Morales A, Jiménez-Díaz L, Platero AI, Bardallo-Pérez J, Díaz-Romero A, Acemel RD, Illán JM, Jiménez-López J, Govantes F. 2016. Biofilm formation-defective mutants in *Pseudomonas putida*. FEMS Microbiol Lett 363:fnw127.

49. Zhang L, Shi Y, Wu Z, Tan G. 2018. Characterization of response regulator GacA involved in phaseolotoxin production, hypersensitive response and cellular processes in *Pseudomonas syringae* pv. *actinidiae* A18. Physiological and Molecular Plant Pathology 103:137–142.

50. Takeuchi K. 2018. GABA, A primary metabolite controlled by the Gac/Rsm regulatory pathway, favors a planktonic over a biofilm lifestyle in *Pseudomonas protegens* CHA0. MPMI 31:274–282.

51. Marutani M, Taguchi F, Ogawa Y, Hossain MdM, Inagaki Y, Toyoda K, Shiraishi T, Ichinose Y. 2008. Gac two-component system in *Pseudomonas syringae* pv. *tabaci* is required for virulence but not for hypersensitive reaction. Mol Genet Genomics 279:313–322.

52. Kidarsa TA, Shaffer BT, Goebel NC, Roberts DP, Buyer JS, Johnson A, Kobayashi DY, Zabriskie TM, Paulsen I, Loper JE. 2013. Genes expressed by the biological control bacterium *Pseudomonas protegens* Pf-5 on seed surfaces under the control of the global regulators GacA and RpoS. Environmental Microbiology 15:716–735.

53. Lalaouna D, Fochesato S, Sanchez L, Schmitt-Kopplin P, Haas D, Heulin T, Achouak W. 2012. Phenotypic switching in *Pseudomonas brassicacearum* involves GacS- and GacA-dependent Rsm Small RNAs. Applied and Environmental Microbiology 78:1658–1665.

54. Martínez-Gil M, Ramos-González MI, Espinosa-Urgel M. 2014. Roles of cyclic di-GMP and the Gac system in transcriptional control of the genes coding for the *Pseudomonas putida* adhesins LapA and LapF. J Bacteriol 196:1484–1495.

55. Driscoll WW, Pepper JW, Pierson LS, Pierson EA. 2011. Spontaneous Gac Mutants of Pseudomonas Biological Control Strains: Cheaters or Mutualists? Applied and Environmental Microbiology 77:7227–7235.

56. Cheng X, de Bruijn I, van der Voort M, Loper JE, Raaijmakers JM. 2013. The Gac regulon of *Pseudomonas fluorescens* SBW25. Environmental Microbiology Reports 5:608–619.

57. Luo Y, Srinivas A, Guidry C, Bull C, Haney CH, Hamilton C. 2025. GacA regulates symbiosis and mediates lifestyle transitions in *Pseudomonas*. mSphere 10:e00277–25.

58. Kim JS, Kim YH, Park JY, Anderson AJ, Kim YC. 2014. The global regulator GacS regulates biofilm formation in *Pseudomonas chlororaphis* O6 differently with carbon source. Can J Microbiol 60:133–138.

59. Li J, Yang Y, Dubern J-F, Li H, Halliday N, Chernin L, Gao K, Cámara M, Liu X. 2015. Regulation of GacA in *Pseudomonas chlororaphis* strains shows a niche specificity. PLOS ONE 10:e0137553.

60. Natsch A, Keel C, Pfirter HA, Haas D, Défago G. 1994. Contribution of the global regulator gene *gacA* to persistence and dissemination of *Pseudomonas fluorescens* biocontrol strain CHA0 introduced into soil microcosms. Applied and Environmental Microbiology 60:2553–2560.

61. Zuber S, Carruthers F, Keel C, Mattart A, Blumer C, Pessi G, Gigot-Bonnefoy C, Schnider-Keel U, Heeb S, Reimmann C, Haas D. 2003. GacS sensor domains pertinent to the regulation of exoproduct formation and to the biocontrol potential of *Pseudomonas fluorescens* CHA0. MPMI 16:634–644.

62. Kim JS, Kim YH, Park JY, Anderson AJ, Kim YC. 2014. The global regulator GacS regulates biofilm formation in *Pseudomonas chlororaphis* O6 differently with carbon source. Can J Microbiol 60:133–138.

63. Song C, Kidarsa TA, van de Mortel JE, Loper JE, Raaijmakers JM. 2016. Living on the edge: Emergence of spontaneous *gac* mutations in *Pseudomonas protegens* during swarming motility. Environmental Microbiology 18:3453–3465.

64. Li E, Zhang H, Jiang H, Pieterse CMJ, Jousset A, Bakker PAHM, de Jonge R. 2021. Experimental-evolution-driven identification of *Arabidopsis* rhizosphere competence genes in *Pseudomonas protegens*. mBio 12:10.1128/mbio.00927-21.

65. Martínez-Granero F, Rivilla R, Martín M. 2006. Rhizosphere selection of highly motile phenotypic variants of *Pseudomonas fluorescens* with enhanced competitive colonization ability. Applied and Environmental Microbiology 72:3429–3434.

66. Chancey ST, Wood DW, Pierson EA, Pierson LS. 2002. Survival of GacS/GacA mutants of the biological control bacterium *Pseudomonas aureofaciens* 30-84 in the wheat rhizosphere. Applied and Environmental Microbiology 68:3308–3314.

67. Ishizawa H, Kuroda M, Inoue D, Ike M. 2022. Genome-wide identification of bacterial colonization and fitness determinants on the floating macrophyte, duckweed. Commun Biol 5:68.

68. Li J, Zhang Y, Jiang W, Zhang L-Q. 2025. Experimental evolution of plant rhizobacteria reveals emerging adaptive mutations. mBio 16:e01023–25.

69. O’Banion BS, Jones P, Demetros AA, Kelley BR, Knoor LH, Wagner AS, Chen J-G, Muchero W, Reynolds TB, Jacobson D, Lebeis SL. 2023. Plant myo-inositol transport influences bacterial colonization phenotypes. Current Biology 33:3111–3124.e5.

70. Schultz D, Wolynes PG, Jacob EB, Onuchic JN. 2009. Deciding fate in adverse times: Sporulation and competence in *Bacillus subtilis*. Proceedings of the National Academy of Sciences 106:21027–21034.

71. O’Malley MR, Dearing HN, Zheng X, Kretschmer AN, Cho TH, Raivio TL, Parsek MR. 2026. The *Pseudomonas aeruginosa* Cpx system provides a cyclic-di-GMP independent link between cell envelope stress and surface sensing. mBio 17.1: e02726–25.

72. Mao M, He L, Yan Q. 2025. An updated overview on the bacterial PhoP/PhoQ two-component signal transduction system. Front Cell Infect Microbiol 15:1509037.

73. Ali-Ahmad A, Fadel F, Sebban-Kreuzer C, Ba M, Pélissier GD, Bornet O, Guerlesquin F, Bourne Y, Bordi C, Vincent F. 2017. Structural and functional insights into the periplasmic detector domain of the GacS histidine kinase controlling biofilm formation in *Pseudomonas aeruginosa*. Sci Rep 7:11262.

74. Liang F, Zhang B, Yang Q, Zhang Y, Zheng D, Zhang L, Yan Q, Wu X. 2020. Cyclic-di-GMP Regulates the Quorum-Sensing System and Biocontrol Activity of Pseudomonas fluorescens 2P24 through the RsmA and RsmE Proteins. Applied and Environmental Microbiology 86:e02016–20.

75. Zhang Y, Zhang B, Wu H, Wu X, Yan Q, Zhang L-Q. 2020. Pleiotropic effects of RsmA and RsmE proteins in *Pseudomonas fluorescens* 2P24. BMC Microbiol 20:191.

76. Collins AJ, Pastora AB, Smith TJ, O’Toole GA. 2020. MapA, a Second Large RTX adhesin conserved across the *Pseudomonads*, contributes to biofilm formation by *Pseudomonas fluorescens*. J Bacteriol 202:10.1128/jb.00277-20.

77. Pastora AB, O’Toole GA. The regulator FleQ both transcriptionally and post-transcriptionally regulates the level of RTX adhesins of *Pseudomonas fluorescens*. J Bacteriol 205:e00152–23.

78. Martínez-Granero F, Navazo A, Barahona E, Redondo-Nieto M, Rivilla R, Martín M. 2012. The Gac-Rsm and SadB signal transduction pathways converge on AlgU to downregulate motility in *Pseudomonas fluorescens*. PLOS ONE 7:e31765.

79. Brinkley DM, Bertolli SK, Gallagher LA, Tan Y, Silva MM de, Brockman A, Zhang D, Peterson SB, Mougous JD. 2025. *Pseudomonads* coordinate innate defense against viruses and bacteria with a single regulatory system. Cell Host & Microbe 33:1333–1346.e7.

80. Liu C, Shi R, Jensen MS, Zhu J, Liu J, Liu X, Sun D, Liu W. 2024. The global regulation of c-di-GMP and cAMP in bacteria. mLife 3:42–56.

81. Azarbad H, Junker RR. 2024. Biological and experimental factors that define the effectiveness of microbial inoculation on plant traits: a meta-analysis. ISME Commun 4:ycae122.

82. Lopes MJ dos S, Dias-Filho MB, Gurgel ESC. 2021. Successful plant growth-promoting microbes: Inoculation methods and abiotic factors. Front Sustain Food Syst 5.

83. Gheorghita AA, Wolfram F, Whitfield GB, Jacobs HM, Pfoh R, Wong SSY, Guitor AK, Goodyear MC, Berezuk AM, Khursigara CM, Parsek MR, Howell PL. 2022. The *Pseudomonas aeruginosa* homeostasis enzyme *AlgL* clears the periplasmic space of accumulated alginate during polymer biosynthesis. Journal of Biological Chemistry 298.

84. Wang Z, Xiong G, Lutz F. 1995. Site-specific integration of the phage ΦCTX genome into the *Pseudomonas aeruginosa* chromosome: characterization of the functional integrase gene located close to and upstream of *attP*. Molec Gen Genet 246:72–79.

85. Morcillo RJL, Zhao A, Tamayo-Navarrete MI, García-Garrido JM, Macho AP. 2020. Tomato root transformation followed by inoculation with *Ralstonia Solanacearum* for straightforward genetic analysis of bacterial wilt disease. JoVE 60302.

86. Hershey DM, Fiebig A, Crosson S. 2021. Flagellar perturbations activate adhesion through two distinct pathways in *Caulobacter crescentus*. mBio 12:e03266–20.

